# Structure-based mechanistic principles underlying the paradoxical effects of pathological RET mutations

**DOI:** 10.1101/2024.12.26.630356

**Authors:** Anna Fassler Bakhman, Michal Cohen, Rachel Kolodny, Mickey Kosloff

## Abstract

The RET receptor is a transmembrane protein that belongs to the receptor tyrosine kinase family. RET activates signaling pathways regulating cell growth, differentiation, and survival in diverse tissues that include the thyroid and enteric nervous system. Mutations that reduce RET levels or disrupt function can cause Hirschsprung’s disease (HSCR), characterized by abnormal distal colon innervation in the developing embryo. In contrast, mutations that constitutively activate RET can lead to tumorigenesis, most notably in multiple endocrine neoplasia type 2 (MEN2) syndromes usually involving the thyroid. Paradoxically, some RET mutations can both reduce RET activity in enteric ganglia, and increase signaling in other tissues, resulting in patients suffering from both HSCR and MEN2A. Although extensive research has been conducted on RET mutations, the structural and mechanistic bases underlying these paradoxical effects remain unclear. Here, we curated data on 70 positions in RET extracellular domains where point-mutations were associated with HSCR, MEN2A, or both. Taking a structure-based approach, we predict the potential effects of mutations in these positions on RET structure. Our analysis suggests that approximately 90% of positions associated with HSCR can, upon mutation, disrupt intramolecular interactions that stabilize RET tertiary structure: residues buried in the protein core, calcium-binding sites, or residues participating in stabilizing intramolecular electrostatic/covalent bonds. A smaller subset of mutations involves substitutions to/from glycines or prolines in key positions. Only a small minority of HSCR-associated positions affect protein-protein interactions needed for signal activation. On the other hand, our analysis showed that ∼75% of mutations in positions that cause MEN2A lead to an unpaired cysteine that can form an intermolecular disulfide bond between two RET monomers. Other mutations that cause MEN2A are also predicted to enhance RET homodimerization via extracellular domains that are proximal to the membrane. Importantly, substitutions that concurrently destabilizes RET tertiary structure and lead to an unpaired cysteine are predicted to cause the paradoxical co-occurrence of HSCR and MEN2A. Our findings suggest a mechanistic basis for almost all identified pathological mutations in RET and imply that therapeutic strategies for targeting RET activity in HSCR and MEN2A may need to be orthogonal.

## INTRODUCTION

The receptor tyrosine kinase *RET* gene encodes for a transmembrane protein that belongs to the receptor tyrosine kinase family. RET initiates signaling pathways that regulate cell growth, differentiation, and survival in diverse tissues that include the embryonic kidney, thyroid, and enteric nervous system (1-3). Abnormal RET signaling can lead to disease, including Hirschsprung’s disease (HSCR) – a congenital malformation characterized by segmental enteric agangliosis, which is often diagnosed in newborns and infants (4-6). Conversely, sustained activation of RET signaling contributes to tumorigenesis and in particular to multiple endocrine neoplasia type 2 (MEN2) syndromes that typically present later in life (7-10). MEN2A, the most common type of MEN2, is characterized in the vast majority of affected individuals by the development of medullary thyroid carcinoma, often presenting in the first to third decade of life. Additional manifestations of MEN2A can include pheochromocytoma and primary hyperparathyroidism, while a variant with cutaneous lichen amyloidosis has also been identified (11, 12). Notably, genetic abnormalities in RET have also been associated with conditions other than HSCR and MEN2, including papillary thyroid carcinoma, non-small cell lung cancer, and congenital anomalies of the urinary tract. However, these RET abnormalities typically involve only the intracellular domains of the protein (13-17).

The RET receptor activates downstream signaling cascades via PI3K/AKT, RAS/RAF/MEK/ERK, JAK2/STAT3, and PLCγ (10). RET activation is typically initiated by binding of any of the following four ligands: Glial cell line-derived growth factor (GDNF), Neurturin (NRTN), Artemin (ARTN), or Persephin – usually combined with binding to one of four GFRa co-receptors (GFRa1-4) (18-20). X-ray structures of RET, solved in complex with these ligands and co-receptors, demonstrated that the extracellular part of the RET receptor is composed of four (D1-D4) cadherin-like domains, followed by one cysteine rich domain (CRD) designated D5 (9, 21, 22) (Fig. 1). In the tetrameric complex of RET with GFRα2 and NTRN, all five extracellular RET domains interact with either ligands or co-receptors (9, 21), while in the dimeric complexes of RET with ligands and co-receptors only four domains (D1-D3, D5) contribute to these interactions (22). The structure of the tetrameric complex further showed that the hexameric 2:2:2 NRTN/GFRα2/RET complex can dimerize, forming a 4:4:4 complex that suppresses RET endocytosis. This suggested that the RET receptor has two distinct interfaces with ligands and two distinct interfaces with its co-receptors.

**Figure 1.**
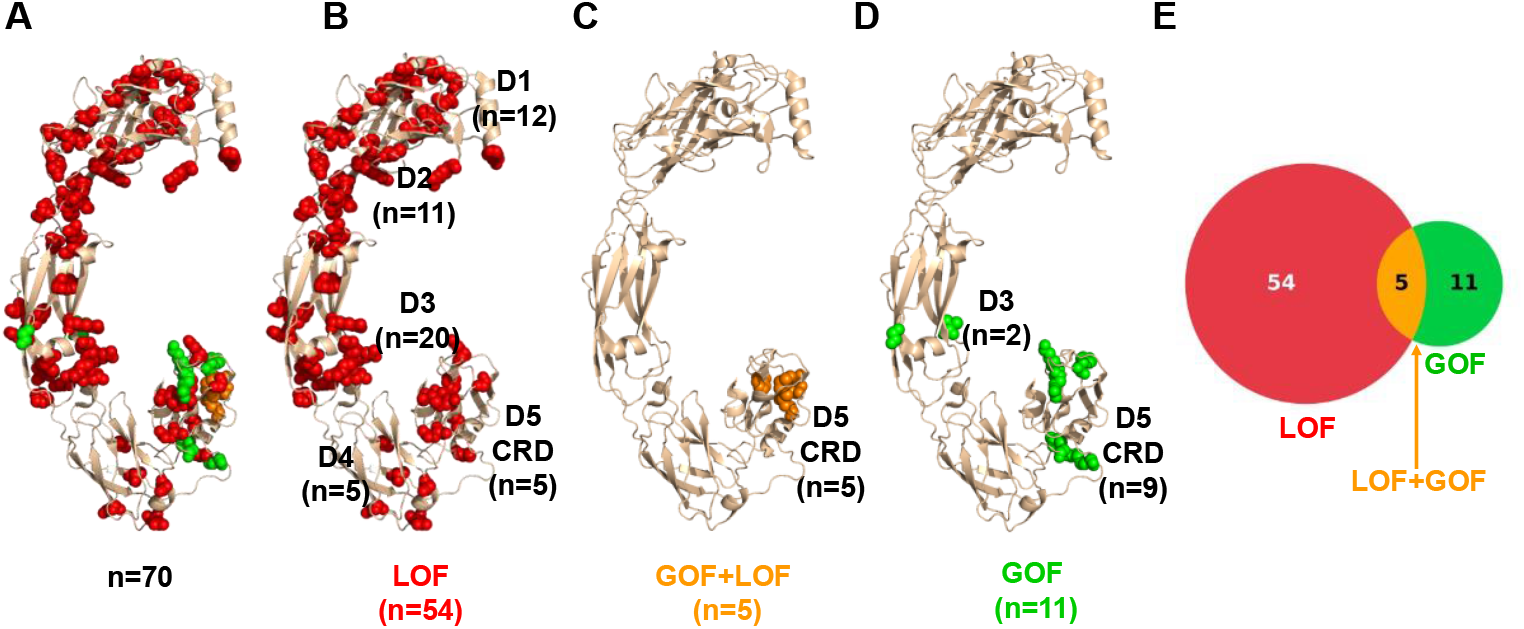
Positions in the extracellular side of RET that are associated with HSRC and/or MEN2A. **A**. GOF and LOF mutations across 70 RET positions that are associated with HSRC and MEN2A, respectively. Positions that are linked only with HSRC (LOF) are shown as red spheres, positions linked only to MEN2A (GOF) are shown as green spheres, and positions associated with both diseases are shown as orange spheres. **B**. LOF positions that are associated with HSRC only, colored as in A. **C**. “Janus” positions that are associated with both HSRC (LOF) and MEN2A (GOF), colored as in A. **D**. GOF positions associated only with MEN2A, colored as in A. **E**. A Venn diagram showing the number of the LOF- and GOF-associated positions shown in panels A-D.

The clinical relevance of the RET extracellular region was demonstrated across studies that mapped gain of function (GOF) mutations in cysteine residues located in the RET CRD leading to MEN2 syndromes by triggering RET dimerization without requiring ligand and co-receptor binding (23-25). Such cysteine mutations presumably result in an unpaired cysteine that induces ligand-independent dimerization and constitutive activation of RET by forming an inter-chain disulfide bond (9, 23, 25). In contrast, loss of function (LOF) mutations in the RET extracellular region can lead to HSCR disease, presumably by affecting the stability of the RET protein – thus reducing functional enteric ganglia in the gut tissue (6, 9, 26-28). Paradoxically, four RET “Janus mutations” in residues C609, C611, C618, and C620 were shown to lead to both LOF and GOF effects and the co-occurrence of both HSCR and MEN2A in the same individual or family (6, 29-36).

In this study, we computationally analyzed structures of the extracellular regions of RET, seeking mechanistic explanations for missense mutation that lead to HSCR, MEN2A, or to the paradoxical occurrence of both diseases. To this end, we identified which positions may perturb RET stability or protein-protein interactions upon mutation and which positions might form intermolecular bonds that can lead to constitutive activation. Our findings map 70 RET positions reported to lead to clinical phenotypes of HSCR or MEN2A and predict the potential structural effects of mutations in all 594 positions that span RET extracellular regions. These insights suggest potential therapeutic avenues for particular mutations, such as stabilizing RET structure for HSCR-associated positions or inhibiting aberrant dimerization with covalent inhibitors to specific cysteine residues for some MEN2A-associated positions.

## RESULTS

### Dataset of RET mutations that lead to diseases

We collected a dataset of 70 positions in the extracellular region of RET in which missense mutations were documented as linked with HSRC and/or MEN2A (Table 1). To this end, we searched the Clinvar database (37) for “pathogenic” and “likely pathogenic” RET variants associated with HSRC disease or MEN2A and found 19 HSRC-associated RET positions and seven MEN2A-associated positions. One of these positions, F555, is associated with both diseases. A search in the COSMIC and cBioPortal databases did not identify additional positions of pathological relevance. We also searched the literature for additional RET positions reported to lead to HSRC or MEN2A (5, 31, 38-42). When the pathological classification of specific positions conflicted between Clinvar and the literature, we prioritized the latter. Specifically, some substitutions in positions C609, C611, C618, and C620 are classified in Clinvar as only GOF mutations (i.e. associated only with MEN2A), while the literature reports the same substitutions as leading to both HSRC and MEN2A (31, 32, 34, 43). Of the 70 RET positions we collected, 54 were linked exclusively to HSCR disease, 11 were associated only with MEN2A, and five were linked to both HSCR and MEN2A.

**Table 1.**
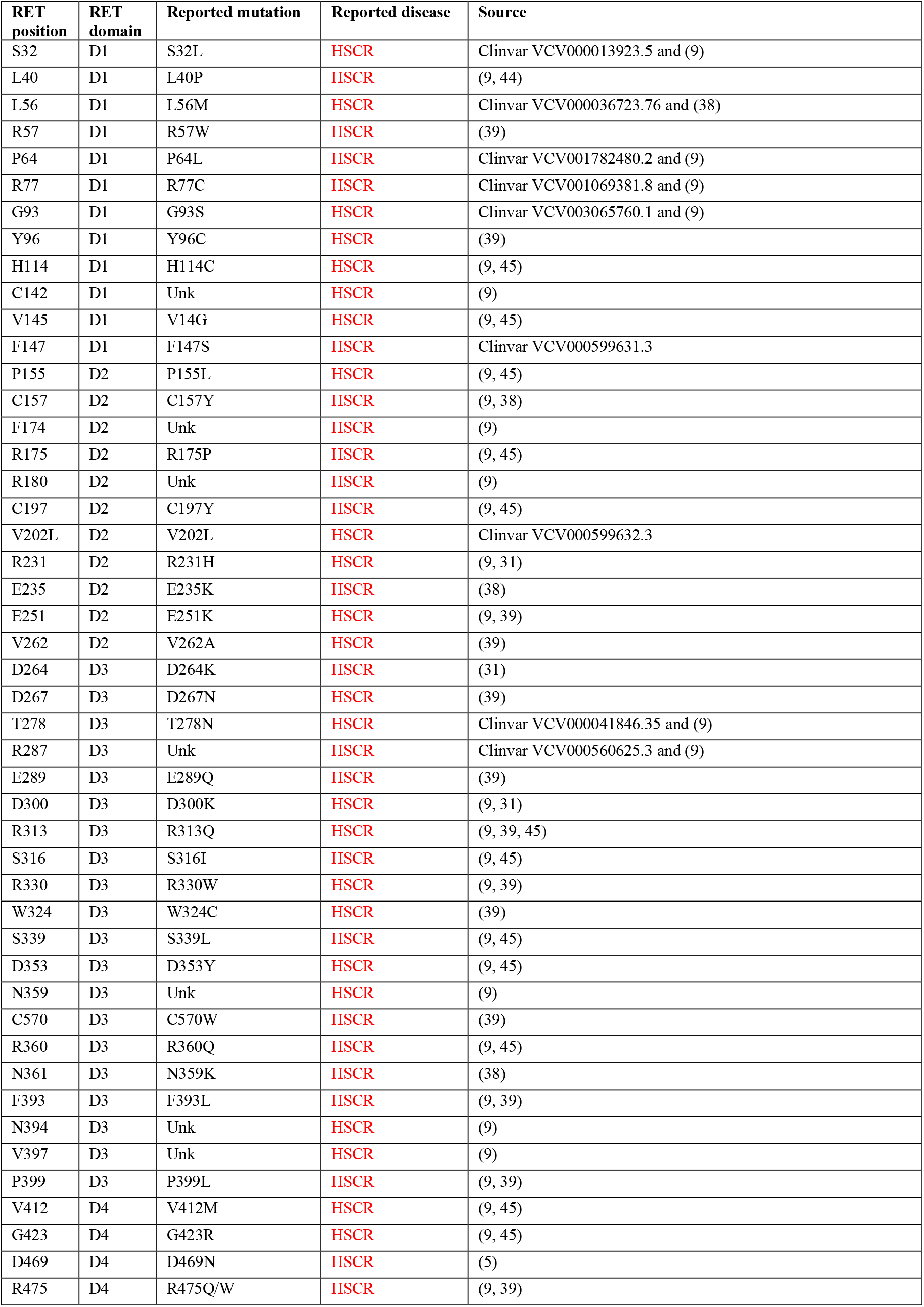
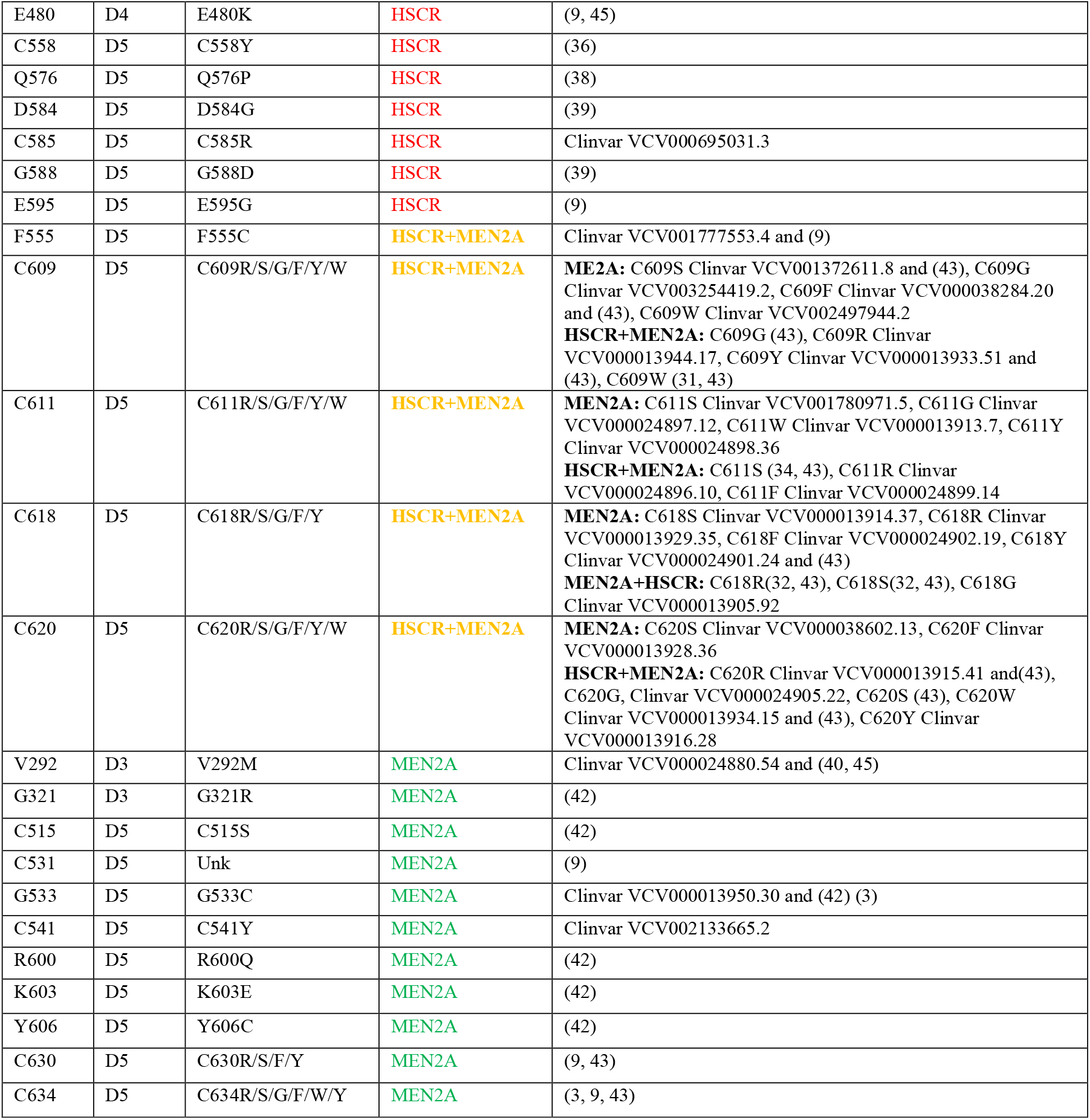
RET missense mutations that lead to HSCR and/or MEN2A. 70 RET positions whose mutations are linked to HSCR, MEN2A, or both. Mutations to unknown amino acids are marked “Unk”.

| RET position | RET domain | Reported mutation | Reported disease | Source |
| --- | --- | --- | --- | --- |
| S32 | D1 | S32L | HSCR | Clinvar VCV000013923.5 and (9) |
| L40 | D1 | L40P | HSCR | (9, 44) |
| L56 | D1 | L56M | HSCR | Clinvar VCV000036723.76 and (38) |
| R57 | D1 | R57W | HSCR | (39) |
| P64 | D1 | P64L | HSCR | Clinvar VCV001782480.2 and (9) |
| R77 | D1 | R77C | HSCR | Clinvar VCV001069381.8 and (9) |
| G93 | D1 | G93S | HSCR | Clinvar VCV003065760.1 and (9) |
| Y96 | D1 | Y96C | HSCR | (39) |
| H114 | D1 | H114C | HSCR | (9, 45) |
| C142 | D1 | Unk | HSCR | (9) |
| V145 | D1 | V14G | HSCR | (9, 45) |
| F147 | D1 | F147S | HSCR | Clinvar VCV000599631.3 |
| P155 | D2 | P155L | HSCR | (9, 45) |
| C157 | D2 | C157Y | HSCR | (9, 38) |
| F174 | D2 | Unk | HSCR | (9) |
| R175 | D2 | R175P | HSCR | (9, 45) |
| R180 | D2 | Unk | HSCR | (9) |
| C197 | D2 | C197Y | HSCR | (9, 45) |
| V202L | D2 | V202L | HSCR | Clinvar VCV000599632.3 |
| R231 | D2 | R231H | HSCR | (9, 31) |
| E235 | D2 | E235K | HSCR | (38) |
| E251 | D2 | E251K | HSCR | (9, 39) |
| V262 | D2 | V262A | HSCR | (39) |
| D264 | D3 | D264K | HSCR | (31) |
| D267 | D3 | D267N | HSCR | (39) |
| T278 | D3 | T278N | HSCR | Clinvar VCV000041846.35 and (9) |
| R287 | D3 | Unk | HSCR | Clinvar VCV000560625.3 and (9) |
| E289 | D3 | E289Q | HSCR | (39) |
| D300 | D3 | D300K | HSCR | (9, 31) |
| R313 | D3 | R313Q | HSCR | (9, 39, 45) |
| S316 | D3 | S316I | HSCR | (9, 45) |
| R330 | D3 | R330W | HSCR | (9, 39) |
| W324 | D3 | W324C | HSCR | (39) |
| S339 | D3 | S339L | HSCR | (9, 45) |
| D353 | D3 | D353Y | HSCR | (9, 45) |
| N359 | D3 | Unk | HSCR | (9) |
| C570 | D3 | C570W | HSCR | (39) |
| R360 | D3 | R360Q | HSCR | (9, 45) |
| N361 | D3 | N359K | HSCR | (38) |
| F393 | D3 | F393L | HSCR | (9, 39) |
| N394 | D3 | Unk | HSCR | (9) |
| V397 | D3 | Unk | HSCR | (9) |
| P399 | D3 | P399L | HSCR | (9, 39) |
| V412 | D4 | V412M | HSCR | (9, 45) |
| G423 | D4 | G423R | HSCR | (9, 45) |
| D469 | D4 | D469N | HSCR | (5) |
| R475 | D4 | R475Q/W | HSCR | (9, 39) |
| E480 | D4 | E480K | HSCR | (9, 45) |
| C558 | D5 | C558Y | HSCR | (36) |
| Q576 | D5 | Q576P | HSCR | (38) |
| D584 | D5 | D584G | HSCR | (39) |
| C585 | D5 | C585R | HSCR | Clinvar VCV000695031.3 |
| G588 | D5 | G588D | HSCR | (39) |
| E595 | D5 | E595G | HSCR | (9) |
| F555 | D5 | F555C | HSCR+MEN2A | Clinvar VCV001777553.4 and (9) |
| C609 | D5 | C609R/S/G/F/Y/W | HSCR+MEN2A | **ME2A:** C609S Clinvar VCV001372611.8 and (43), C609G Clinvar VCV003254419.2, C609F Clinvar VCV000038284.20 and (43), C609W Clinvar VCV002497944.2<br>**HSCR+MEN2A:** C609G (43), C609R Clinvar VCV000013944.17, C609Y Clinvar VCV000013933.51 and (43), C609W (31, 43) |
| C611 | D5 | C611R/S/G/F/Y/W | HSCR+MEN2A | **MEN2A:** C611S Clinvar VCV001780971.5, C611G Clinvar VCV000024897.12, C611W Clinvar VCV000013913.7, C611Y Clinvar VCV000024898.36<br>**HSCR+MEN2A:** C611S (34, 43), C611R Clinvar VCV000024896.10, C611F Clinvar VCV000024899.14 |
| C618 | D5 | C618R/S/G/F/Y | HSCR+MEN2A | **MEN2A:** C618S Clinvar VCV000013914.37, C618R Clinvar VCV000013929.35, C618F Clinvar VCV000024902.19, C618Y Clinvar VCV000024901.24 and (43)<br>**MEN2A+HSCR:** C618R(32, 43), C618S(32, 43), C618G Clinvar VCV000013905.92 |
| C620 | D5 | C620R/S/G/F/Y/W | HSCR+MEN2A | **MEN2A:** C620S Clinvar VCV000038602.13, C620F Clinvar VCV000013928.36<br>**HSCR+MEN2A:** C620R Clinvar VCV000013915.41 and(43), C620G, Clinvar VCV000024905.22, C620S (43), C620W Clinvar VCV000013934.15 and (43), C620Y Clinvar VCV000013916.28 |
| V292 | D3 | V292M | MEN2A | Clinvar VCV000024880.54 and (40, 45) |
| G321 | D3 | G321R | MEN2A | (42) |
| C515 | D5 | C515S | MEN2A | (42) |
| C531 | D5 | Unk | MEN2A | (9) |
| G533 | D5 | G533C | MEN2A | Clinvar VCV000013950.30 and (42) (3) |
| C541 | D5 | C541Y | MEN2A | Clinvar VCV002133665.2 |
| R600 | D5 | R600Q | MEN2A | (42) |
| K603 | D5 | K603E | MEN2A | (42) |
| Y606 | D5 | Y606C | MEN2A | (42) |
| C630 | D5 | C630R/S/F/Y | MEN2A | (9, 43) |
| C634 | D5 | C634R/S/G/F/W/Y | MEN2A | (3, 9, 43) |

We mapped these 70 positions onto the 3D structure of the extracellular region of RET – classified as LOF, GOF, or both (Fig. 1A). The 59 positions associated with LOF mutations are distributed across all five domains, but with fewer positions in D4 and in the CRD D5 (Fig. 1B). Notably, the five RET mutations that can lead to both diseases are only in D5 (Fig. 1C) while GOF-only mutations are predominantly in the CRD D5 – 14 out of 16 (Fig. 1D). We note that almost all of the 16 GOF mutations (Fig. 1E) seem to be clustered in positions located in a specific region of CRD D5 (Fig. 1C, D).

### Predicted mechanistic principles for LOF effects in RET positions associated with HSCR

We hypothesize that mutations that reduce downstream signaling by RET or that destabilize the protein and thereby reduce its amount in cells can lead to RET LOF and thereby to HSCR disease. Mutations in RET positions involved in protein-protein interactions with co-receptors or ligands may impair RET activation and reduce downstream signaling. Mutations that, by diverse mechanisms, destabilize RET, can also lead to LOF. One such mechanism may involve calcium-binding, as calcium ions were shown to stabilize RET tertiary structure (9). Therefore, mutations in positions involved in calcium-binding may destabilize the receptor and disrupt its folding. Alternatively, substitutions of RET positions that are part of its protein core and stabilize its tertiary structure could also disrupt folding or structural integrity. Lastly, mutations in RET that disrupt strong intramolecular interactions may also affect its tertiary structure. Therefore, we mapped which residues mediate interactions between RET and its co-receptors and ligands, participate in the calcium-binding sites, are buried in the protein core, introduce a specific disruption of RET structures via amino acids with unique physico-chemical properties such as glycines or prolines, or may affect RET stability by disrupting strong intramolecular interactions (Supplementary Fig. 1, Fig. 2).

We examined these 70 positions for effects on RET protein-protein interactions by analyzing the experimental structures of RET in complex with each of the following four co-receptors and four ligands: RET with GFRα1 and GDNF (PDB ID 6Q2J), RET with GFRα3 and Artemin (PDB ID 6Q2S), RET with GFRAL and GDF15 (PDB ID 6Q2N), RET with GFRα2 and Neurturin (PDB ID 6Q2O), and the 4:4:4 structure of RET with GFRα2 and Neurturin (PDB ID 6Q2R) (9). RET residues located within 5 Å of the interface with co-receptors (Fig. 2A) or ligands (Fig. 2B) were classified as “interacting residues”. We identified 43 RET residues within 5Å of a co-receptor and 24 RET residues within 5Å of a ligand (Supplementary Fig. 2, Fig. 2A,B). RET residues that interact with co-receptors are mostly in domains D1 and D2, with only a few residues in D5. In contrast, RET residues that interact with ligands are only in domains D4 and D5. Among the 59 HSCR-associated RET positions that presumably reduce activity upon mutation, only four are in proximity to RET co-receptors, R77, H114, R175, and D353 (Fig. 2A) and only one position associated with HSCR, E595, is within 5Å of a ligand (Fig. 2B).

**Figure 2.**
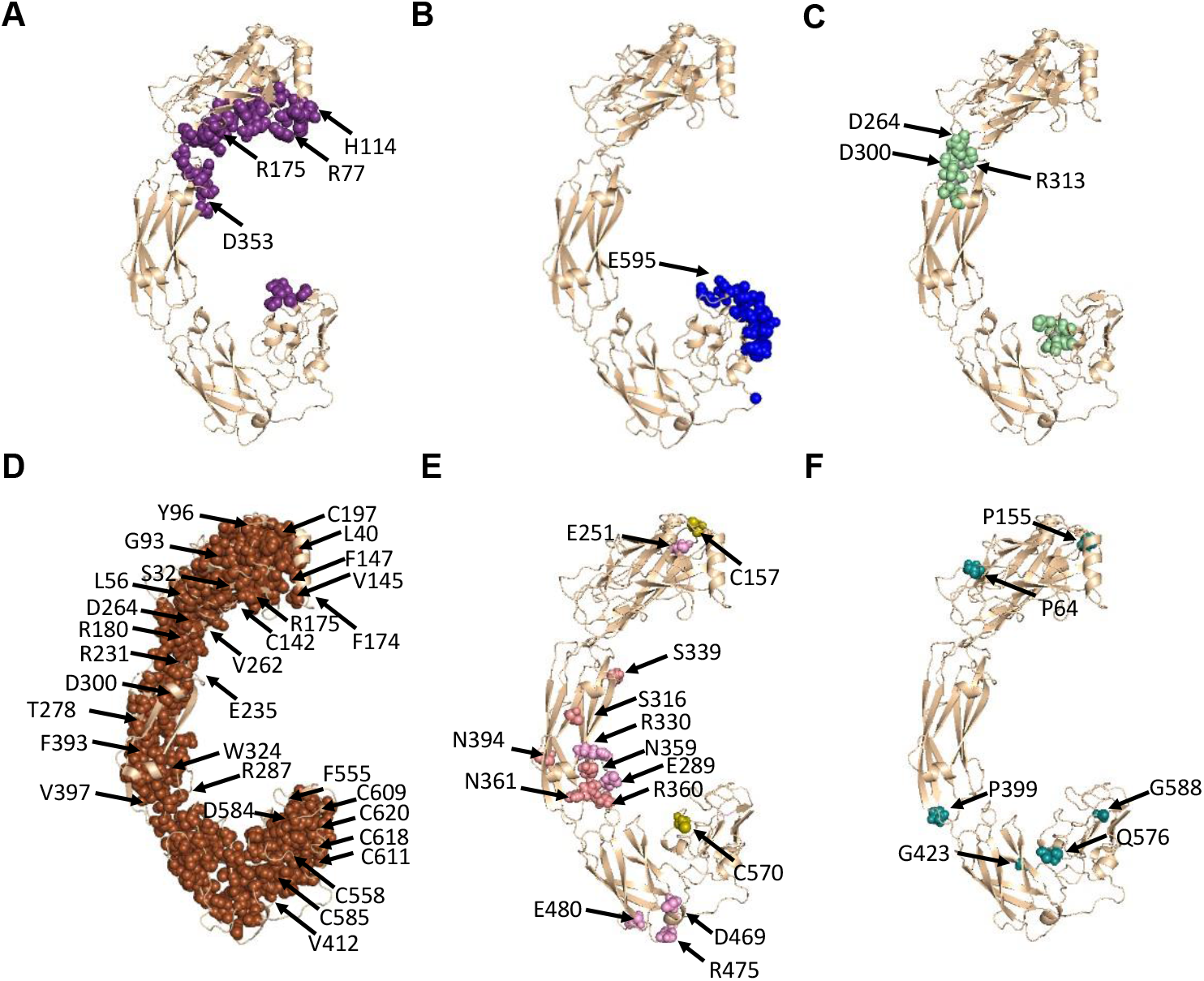
Predicted structural basis for RET LOF mutations leading to HSCR disease. **A**. 43 RET residues predicted to interact with RET co-receptors, shown as purple spheres. The four mutated positions among these residues that lead to HSCR are labeled with black arrows. **B**. 24 RET residues predicted to interact with RET ligands, shown as blue spheres. The one mutated position that leads to HSCR is labeled with a black arrow. **C**. 19 RET residues that bind calcium ions, shown as light green spheres. The three mutated positions among these residues that lead to HSCR are labeled with black arrows. **D**. 222 RET residues that are classified as “buried” in the protein core (rASA ≤ 5% or BSA ≥ 200 Å^2^), shown as brown spheres. The 31 mutated positions among these residues that lead to HSCR are labeled with black arrows. **E**. 14 RET residues that lead to HSCR and stabilize RET tertiary structure via strong intramolecular interactions, labeled with black arrows and shown as spheres colored light pink (participating in hydrogen bonds), pink (participating in salt bridges), and olive (forming intermolecular disulfide bonds). **F**. Six RET positions that involve mutations to/from prolines/glycines, thereby destabilizing RET tertiary structure, labeled with black arrows and shown as teal spheres.

We next mapped positions involved in calcium-binding (Fig. 2C), which was shown to be important for RET stability (9). Out of the 22 RET positions predicted to be involved in stabilizing the tertiary structure of RET through calcium-binding, two positions, D264 and D300, were associated with HSCR. A third HSCR-associated residue, R313, is immediately adjacent to the calcium-binding Y314, and was therefore also classified as part of the calcium binding site. Therefore, three of the 59 LOF RET positions participate in calcium binding and likely lead to HSCR through destabilization of RET tertiary structure.

To map RET residues that are part of the core of the protein, we identified positions that are sufficiently buried in the protein hydrophobic core, where a mutation is more likely to perturb RET folding or destabilize the structure. To quantify the burial of each RET residue, we used an approach developed in our previous study (46). We measured the accessible surface area (ASA) of each RET residue using surfv (47). Then, we calculated the relative accessible surface area (rASA) for each residue by dividing the ASA value by the maximal empirical ASA for each residue, taken from (48). We also calculated the buried surface area (BSA) for each residue – as the difference between the maximal empirical ASA value and the calculated ASA for each residue. We classified RET residues as “buried” when either their rASA values are ≤15% or their BSA values are > 200Å^2^, with all other residues considered “exposed”. This burial-based analysis classified the 594 residues in the RET extracellular region as follows – 222 residues as “buried” and 372 as “exposed” (Fig. 2D). We predict that these 222 buried residues are more likely to destabilize RET tertiary structure upon mutations that change the physico-chemical properties or size of the residue and thereby decrease RET levels in the relevant tissues. Indeed, 31 out of 59 RET LOF positions involve buried residues. We also note that two calcium-binding residues and one co-receptor binding residue are also classified as buried, meaning that a mutation of these residues can lead to LOF due to multiple simultaneous causes. Taken together, 32 of the 59 LOF mutations in RET destabilize its tertiary structure by affecting the protein core and/or calcium-binding, and only four residues reduce its activity solely via protein-protein interactions with ligands or co-receptors. This leaves 23 RET positions that were not classified under the categories above.

14 out of these 23 RET positions associated with HSCR participate in intramolecular interactions that can affect RET stability and folding (Fig. 2E). Visual inspection identified that two of these 14 HSCR-associated RET positions (C157 and C570) are cysteine residues that participate in intramolecular disulfide bonds with residues C197 and C585, respectively. The other 12 RET residues participate in intramolecular hydrogen bonds or salt-bridges. E251, S316, and E480 form intramolecular hydrogen bonds with R205, H333, and T503, respectively. R330 forms an intramolecular salt-bridge network with three negatively charged residues – E289, D290, and E332. Another salt-bridge is between D469 and R475, and two hydrogen bond networks are formed between S339, N359, R360, and N361, and between N394, Q371, and H392.

Six of the remaining nine HSCR-associated RET positions involve substitutions either to or from proline or glycine (Fig. 2F). Such mutations are likely to affect RET stability because of the extreme backbone flexibility or rigidity associated with glycines and prolines, respectively (49, 50). P64 is in the middle of a loop next to a buried residue, F66. Consistent with previous findings (5), we predict that the HSCR-associated P64L mutation may lead to steric interference with F66, thereby affecting RET tertiary structure and stability. Similar to P64, P155 is in a loop and is also located between two buried residues, L160 and Y41. Likewise, P339 is adjacent to the buried residue V400. Mutations in either of these prolines will likely perturb these regions. Reciprocally, Q576, which is also positioned in the middle of a loop in the D4-D5 interface, is substituted by proline in an HSCR-associated mutation, thereby increasing conformational rigidity and likely affecting RET tertiary structure negatively. In contrast, the G423R mutation inserts both a charge and a significant steric bulk next to two large residues, K424 and Q421, thereby likely perturbing RET tertiary structure. We note that five additional residues that were classified above as buried or involved in protein-protein interactions, also involve mutations to or from prolines/glycines. L40P, G93S, and D584P are mutations in buried residues, R175P is a mutation in a residue that is both buried and predicted to bind co-receptors, while E595P is a mutation in a residue predicted to bind ligands. Therefore, these positions can also contribute to HSCR through multiple distinct mechanisms.

Overall, our analysis reveals that in 52 out of the 59 HSCR-associated LOF mutations, the involved RET residues are important to its folding or stabilize its tertiary structure, thus providing mechanistic structural-based explanations to the association of these mutations with LOF/HSCR. Only four mutations appear to contribute to HSCR by impacting solely protein-protein interactions. For only three HSCR-associated positions out of the 59, our analysis did not predict an effect on protein-protein interactions nor on protein stability – R57, V202, and D267. These are “exposed” residues located in D2, D3, and D4, respectively, and are not predicted to interact with ligands or co-receptors. R57 is positioned in the middle of a loop, directly adjacent to proline and arginine residues. It is possible that the mutation of this charged arginine to a bulky tryptophan (Table 1) alters local intra-molecular interactions and thereby global stability. In contrast, V202 and D267 are immediately adjacent to residues that are buried within the protein core. Specifically, V202 is immediately adjacent to the buried residue I200, while D267 is immediately adjacent to D266 and E178, and is also close to E265, all of which are buried yet charged residues. It is therefore plausible that mutations in these two residues will indirectly affect these buried residues and thereby RET tertiary structure and stability.

### Predicted mechanistic principles for RET positions associated with GOF effect and MEN2A

Out of the 16 gain-of-function (GOF) positions identified in RET (Table 1), 12 mutations involve substitutions to or from cysteine residues that participate in disulfide bonds within the RET CRD D5. We hypothesize that these substitutions lead to an unpaired cysteine in D5 that can form an intermolecular disulfide bond with its unpaired counterpart across the RET dimer interface. Such an intermolecular disulfide bond will lead to aberrant RET homodimerization and thus to increased signaling that presumably underlies oncogenesis. Nine of these 12 mutations change a cysteine residue to a non-cysteine residue (Fig. 3A). Interestingly, the remaining three out of 12 GOF mutations, F555C, G533C, and Y606C, also lead to an unpaired cysteine but via a reciprocal route; instead of mutating half of an existing intramolecular disulfide bridge, they introduce a novel, unpaired, cysteine (Fig. 3B).

**Figure 3.**
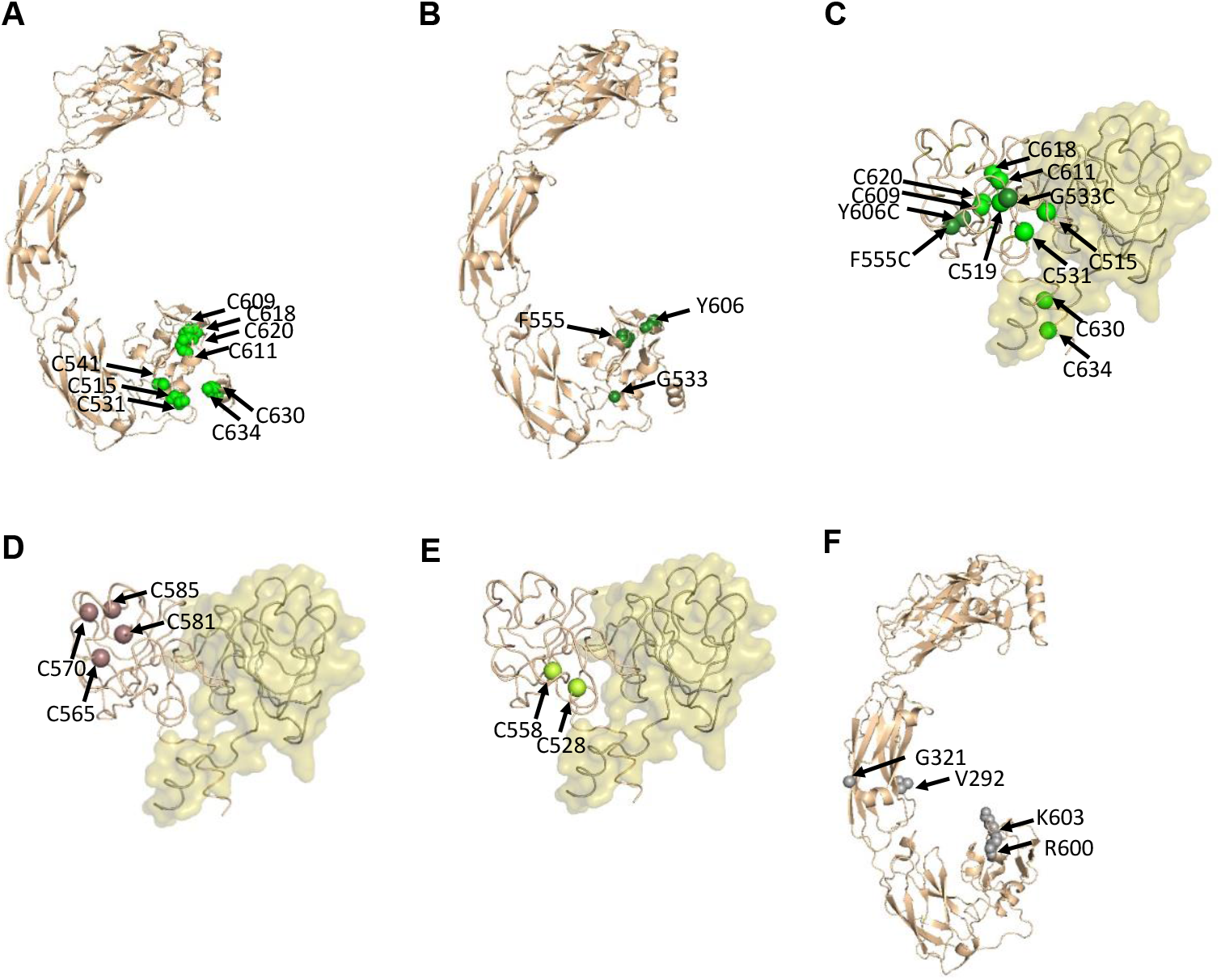
Predicted structural basis for RET GOF mutations that lead to MEN2A. **A**. Nine cysteine residues which, upon mutation to a non-cysteine residue, result in an unpaired cysteine – shown as green spheres. **B**. Three non-cysteine residues which, upon mutation to a cysteine, also result in an unpaired cysteine – shown as dark green spheres. **C**. Positions of the 12 unpaired cysteines resulting from the mutations described in A and B, mapped onto one of the monomers in an Alphafold3 model of the D5-D5 dimer. Cα atoms of these unpaired cysteines are shown as spheres and colored as in A and B. Note that eight of these 12 GOF mutations involve both cysteines across the same intramolecular disulfide bond. **D**. Four cysteine residues in D5 that have not been associated with MEN2A, mapped onto one of the monomers in the model of the D5-D5 dimer (from C), with the Cα atoms shown as dark pink spheres. **E**. Positions of the two cysteine residues in D5 that are close to the predicted D5-D5 interface but have not been associated with MEN2A, with their Cα atoms shown as light green spheres. **F**. Four RET positions that lead to GOF upon mutation but do not involve substitutions to or from cysteines, shown as grey spheres.

In most GOF cases in patients, we find substitutions in either cysteine across the same disulfide bond. However, in the disulfide bond between C541 and C519, only mutations in C541 have been observed, leading to an unpaired C519 (Fig. 3C). All of these unpaired cysteines can potentially form an intermolecular disulfide bond with their counterparts across the dimer interface, as detailed above – similarly leading to RET homodimerization that can be independent of ligands and co-receptors. Indeed, we observed that all of the 12 unpaired cysteines that result from a GOF mutation are clustered around the same outer-facing side of D5 (Fig, 3A, B), supporting the intermolecular dimerization hypothesis. To test this hypothesis, we modeled the D5-D5 dimer of RET using AlphaFold3 (51). This model shows that all 12 unpaired cysteine positions are in the region of D5 that faces the predicted homo-dimerization interface – 11 of the 12 unpaired GOF cysteine residues mentioned above are ≤ 9Å from the modelled dimer interface and F555C is ∼12Å from this interface (Fig. 3C). In contrast, almost all cysteine residues in D5 that have not been associated with MEN2A are farther away from the predicted D5-D5 dimer interface, at distances of 13-18Å (Fig. 3D). The only exception is C528 that is 7Å from the predicted D5-D5 dimer interface (Fig. 3E); this residue forms an intramolecular disulfide bond with C558 but has not been reported to be associated with MEN2A. Therefore, this location suggests that a mutation of C558 may also lead to a GOF and to MEN2A.

The remaining four of the 16 reported GOF mutations do not lead to unpaired cysteines and therefore contribute to MEN2A through a different mechanism (Fig. 3F). These mutations are in four positions that are located in domains D3 (V292M, G321R) and D5 (R600W, K603E). V292 and G321 are exposed residues that are far from both the ligand-binding and co-receptor binding regions. The V292M mutation does not change the physico-chemical properties of this residue sufficiently to explain the GOF phenotype. However, the G321R mutation, as it involves a dramatic change in the physico-chemical properties of the residue, might induce an intramolecular conformational change that can increase RET activation. As for the two residues in D5, both R600 and K603 are close to the predicted D5-D5 dimer interface (Fig. 3C), and mutations of these positions might therefore also promote D5-D5 dimerization, albeit through a non-covalent mechanism that probably requires a conformational change. We note that K603 is within 5Å of the co-receptor, but visual inspection did not identify RET/co-receptor interactions that might be enhanced as a result of this mutation and can explain the GOF phenotype. We also note that the R600 residue, which is also positioned in close proximity to co-receptor binding residues, has been reported to be substituted in extremely rare cases (52), so perhaps the GOF phenotype is less pronounced. Overall, our analysis suggests that 14 of the 16 GOF mutations associated with MEN2A may enhance D5-D5 dimerization and thereby enhance downstream signaling, promoting tumorigenesis.

### Predicted mechanistic explanation for the paradoxical LOF and GOF effects in RET Janus positions associated with both HSCR and MEN2A

Five disease-associated RET positions – F555, C609, C611, C618, and C620 – are linked to both HSCR and MEN2A (Table 1, Fig. 1). Our analysis revealed that all of these positions contribute to the stability of the tertiary structure of RET, as they are part of the protein core (Fig. 1D). Interestingly, it has been documented in patients that F555 is substituted exclusively to cysteine, while C609, C611, C618, and C620 are consistently substituted to the same specific amino acids – arginine, serine, glycine, phenylalanine, tyrosine, or tryptophan (Table 1). Therefore, all five Janus mutations involve substantial changes in physico-chemical properties and are likely to affect protein folding. These substantial substitutions, particularly in such core residues, may destabilize the tertiary structure of RET, impairing the embryonic development of the enteric nervous system and contributing to HSCR. On the other hand, mutations in all five residues, which are all located at the periphery of D5, result in an unpaired cysteine (Fig. 3). F555C introduces a new and unpaired cysteine to the predicted D5-D5 interface, whereas the C609, C611, C618, and C620 positions are all part of intramolecular disulfide bonds, so their mutation leaves an unpaired cysteine close to the predicted D5-D5 interface. These unpaired cysteines can lead to aberrant RET dimerization as detailed above and contribute to MEN2A, which, unlike HSCR, manifests later in life.

## DISCUSSION

We curated a comprehensive dataset of 70 positions in the extracellular domains of RET that have been associated with either HSCR disease, MEN2A, or both – integrating entries from both the Clinvar database and the literature (Tables 1, 2). This dataset provides a reconciled and clarified view of RET mutations that is more comprehensive than previously published and, to our knowledge, is the largest dataset of disease-associated RET mutations. We also provide predicted structural and mechanistic bases for the LOF and GOF phenotypes for almost all of these disease-associated mutations across these 70 RET positions. Our carefully curated dataset can serve as a benchmark for developing structure-informed Variant Effect Predictors (VEPs) (53), which are promising as powerful tools for missense evaluation in the context of human health. Our detailed analysis of both GOF and LOF mutations also provides insights into how one should design such tools. For example, our analysis supports the common wisdom that LOF mutations tend to be broadly distributed, while GOF mutations show more spatial clustering (53). Importantly, the RET example highlights that LOF and GOF mutations are not necessarily mutually exclusive.

**Table 2.** RET missense mutations that lead to HSCR, MEN2A, and both diseases – categorized by their predicted effect or mechanism. Mutations are classified based on intra- and intermolecular interactions, burial status, calcium-binding participation, and the type of residue substitution (such as glycine/proline mutations in key positions or mutation of/to cysteine).

| Hirschsprung |  | MEN2A |
| --- | --- | --- |
| LOF | LOF+GOF | GOF |
| S32<sup>Bur</sup>, L40<sup>Bur</sup>, L56<sup>Bur</sup>, R57, P64<sup>G/P</sup>, R77<sup>CR</sup>, G93<sup>Bur</sup>, Y96<sup>Bur</sup>, H114<sup>CR</sup>, C142<sup>Bur/Cys</sup>, V145<sup>Bur</sup>, F147<sup>Bur</sup>, P155<sup>G/P</sup>, C157<sup>Cys</sup>, F174<sup>Bur</sup>, R175<sup>CR/Bur</sup>, R180<sup>Bur</sup>, C197<sup>Bur/Cys</sup>, V202, R231<sup>Bur</sup>, E235<sup>Bur</sup>, E251<sup>SB</sup>, V262<sup>Bur</sup>, D264<sup>Ca/Bur</sup>, D267, T278<sup>Bur</sup>, R287<sup>Bur</sup>, E289<sup>SB</sup>, D300<sup>Ca/Bur</sup>, R313<sup>Ca</sup>, S316<sup>HB</sup>, W324<sup>Bur</sup>, R330<sup>SB</sup>, S339<sup>HB</sup>, D353<sup>CR</sup>, N359<sup>HB</sup>, R360<sup>HB</sup>, N361<sup>HB</sup>, F393<sup>Bur</sup>, N394<sup>HB</sup>, V397<sup>Bur</sup>, P399<sup>G/P</sup>, V412<sup>Bur</sup>, G423<sup>G/P</sup>, D469<sup>SB</sup>, R475<sup>SB</sup>, E480<sup>HB</sup>, C558<sup>Bur/Cys</sup>, C570<sup>Cys</sup>, Q576<sup>G/P</sup>, D584<sup>Bur</sup>, C585<sup>Bur/Cys</sup>, G588<sup>G/P</sup>, E595<sup>Lig</sup> | F555<sup>Bur/Cys</sup>, C609<sup>Bur/Cys</sup>, C611<sup>Bur/Cys</sup>, C618<sup>Bur/Cys</sup>, C620<sup>Bur/Cys</sup> | V292, G321, C515<sup>Cys</sup>, C531<sup>Bur/Cys</sup>, G533<sup>Bur</sup>, C541<sup>Bur/Cys</sup>, R600, K603, Y606<sup>Cys</sup>, C630<sup>Cys</sup>, C634<sup>Cys</sup> |
| Bur – buried residue<br>G/P – glycine/proline in key positions<br>CR – interactions with co-receptor<br>Ca – calcium binding | HB – intramolecular hydrogen bond<br>SB – intramolecular salt-bridge<br>Lig – interactions with ligand<br>Cys – mutation of/to cysteine |  |

Our structure-based analysis of this dataset suggests that ∼90% of RET extracellular positions associated with HSCR may destabilize the tertiary structure of the RET receptor. Only a small minority of mutations appear to cause LOF by attenuating protein-protein interactions with the co-receptors or ligands that activate RET. A previous analysis of disease-associated missense mutations across dozens of different proteins reached similar ratios, estimating ∼80% of such mutations impair protein stability, with the majority of these involving residues buried in the protein core or causing disruption of strong intramolecular interactions (54). Indeed, our predictions suggest that the most common reason for RET destabilization and therefore LOF is perturbation of the protein core. However, disruption of strong intramolecular interactions that include salt-bridges, hydrogen bonds, and disulfide bonds, or mutations at calcium-binding sites that were shown to stabilize RET tertiary structure, can also lead to LOF and HSCR disease. An additional mechanism for RET destabilization can occur when residues in the protein core or in intramolecular interfacial positions are substituted to or from glycines or prolines, due to the unique effect of such mutations on protein 3D structure. Mutations involving glycine, with its extreme flexibility, or proline, which rigidifies the protein backbone and can adopt unique conformations, can disrupt local folding and global stability. Notably, some HSCR-associated positions can lead to LOF via two or even three different mechanisms. Specifically, mutations in D264 and D300 can destabilize RET not only by disrupting calcium binding, but also by compromising the protein core. Similarly, mutations in the buried residues L40, G93, and E584 can affect both the protein core due to their burial, or through the specific conformational effects of a substitution to or from glycine. Furthermore, the buried residue R175 is predicted to affect interactions with co-receptors but also involves a substitution to proline. Likewise, the E595P mutation is also predicted to affect ligand binding. Overall, we predict that mutations destabilizing the protein tertiary structure are the most likely to lead to HSCR.

In contrast to LOF mutations observed in HSCR, GOF mutations associated with MEN2A predominantly involve mutations to or from cysteine residues in the CRD D5 domain of RET. Such mutations lead to an unpaired cysteine close to the predicted D5-D5 dimer interface that presumably forms an intermolecular disulfide bond across the homodimer. This, in turn, leads to aberrant RET dimerization and activation that is less dependent on co-receptors and ligands. There are four cysteine residues in D5 that are not known to lead to GOF upon mutation, which our analysis suggests is due to their more distant location from the predicted dimer interface – presumably not allowing the formation of an intermolecular disulfide bond across the dimer interface. The additional GOF mutations that do not involve unpaired cysteines are also located in membrane-proximal domains, supporting a common GOF mechanism of enhancing RET homodimerization via these membrane-proximal domains. Two of these positions are located in D5, where they can enhance D5-D5 dimerization. A third position might involve activity-enhancing conformational changes, also supporting a general mechanism of GOF mutations leading to constitutive activation by enhancing RET dimerization proximal to the membrane. Indeed, these membrane-proximal interactions resemble other homotypic protein-protein interactions, and in particular the juxta-membrane homodimerization shown to occur between other RTKs – PDGFR, KIT, and VEGFR receptors (55-57). Finally, we predict that a mutation in C558 can lead to an unpaired C528 and thus contribute to MEN2A – a mutation that has not been identified in patients to date.

Our structural analysis suggests a mechanistic explanation for the seemingly paradoxical five RET Janus positions that lead to both HSCR and MEN2A. Recent work has highlighted the complexity of RET Janus mutations, showing that mice carrying the RET C618F mutation, when combined with a reduction in RET gene dosage, developed intestinal aganglionosis due to aberrant RET activation (58). This, in turn, led to premature neuronal differentiation and impaired precursor migration. Our analysis showed that all Janus positions are buried within D5, and the substitutions found in HSCR involve dramatic changes to the physico-chemical properties of these residues, explaining the LOF phenotype by a substantial destabilization of RET. On the other hand, all five mutations result in an unpaired cysteine that is predicted to form a D5-D5 intermolecular disulfide bond, leading to the GOF phenotype that is explained as detailed above. The concurrent prediction though is that the LOF phenotype is partial, allowing sufficient levels of RET to facilitate abnormal dimer formation and MEN2A later in life. More broadly, the dual pathogenic phenotype of mutations in RET highlights the need for precisely tailored therapeutic strategies to address RET-associate diseases in the thyroid and enteric ganglia. Stabilizing the RET protein might help mitigate the LOF HSCR phenotype in the intestine, while preventing aberrant receptor activation is crucial for treating MEN2A. Furthermore, the pivotal role of cysteine residues within D5 in RET regulation underscores the potential of developing covalent inhibitors targeting intermolecular disulfide bonds as a therapeutic strategy for MEN2A.

## MATERIALS AND METHODS

### Protein structures

We used the following 3D structures in our analysis and visualization of RET with different co-receptors and ligands: RET with GFRα1 and GDNF (PDB ID 6Q2J), RET with GFRα3 and Artemin (PDB ID 6Q2S), RET with GFRAL and GDF15 (PDB ID 6Q2N), RET with GFRα2 and Neurturin (PDB ID 6Q2O), and 4:4:4 structure of RET with GFRα2 and Neurturin (PDB ID 6Q2R). Short segments of RET (residues 129-137, 207-211, 246-251, and 379-387) that are missing in PDB entries 6Q2R, 6Q2O, 6Q2S, and 6Q2S were modeled using Loopy (59) and partial or missing side chains were modeled using Scap (59). We used the Alpahfold3 server (https://alphafoldserver.com/) (51) to predict the 3D structure of the RET D5-D5 dimer and PyMol (https://www.pymol.org/) for structural visualization.

### Structure-based prediction of mutation effect

RET residues ≤ 5 Å of the interface with co-receptors and ligands were classified as “interacting residues”. Calcium-binding residues were classified according to Li *et al*. (9). To map residues that can affect the tertiary structure of a protein due to being buried in the protein core, we followed the methodology described previously (46). We measured the accessible surface area (ASA) of each residue using surfv (47). Then, we calculated the relative accessible surface area (rASA) for each residue by dividing its ASA by the maximal empirical ASA, taken from the values calculated from a large dataset of structures culled from the PDB in (48), who followed the approaches laid out by Rose *et al*. (60) and Miller *et al*. (61). Buried surface area (BSA) was calculated by subtracting the ASA of each residue from the maximal empirical ASA value for that residue. A residue was classified as “buried” when its rASA ≤ 15% or when its BSA ≥ 200 Å^2^. Residues involved in intramolecular salt bridges or hydrogen bonds were identified by visual inspection.

## Notes

### Competing Interest Statement

The authors have declared no competing interest.

### Summary of Updates

Title updated Revised Figures Supplemental files updated.

