## Supplementary Figures 1 and 2 for "Structure-based mechanistic principles underlying the paradoxical effects of pathological RET mutations"

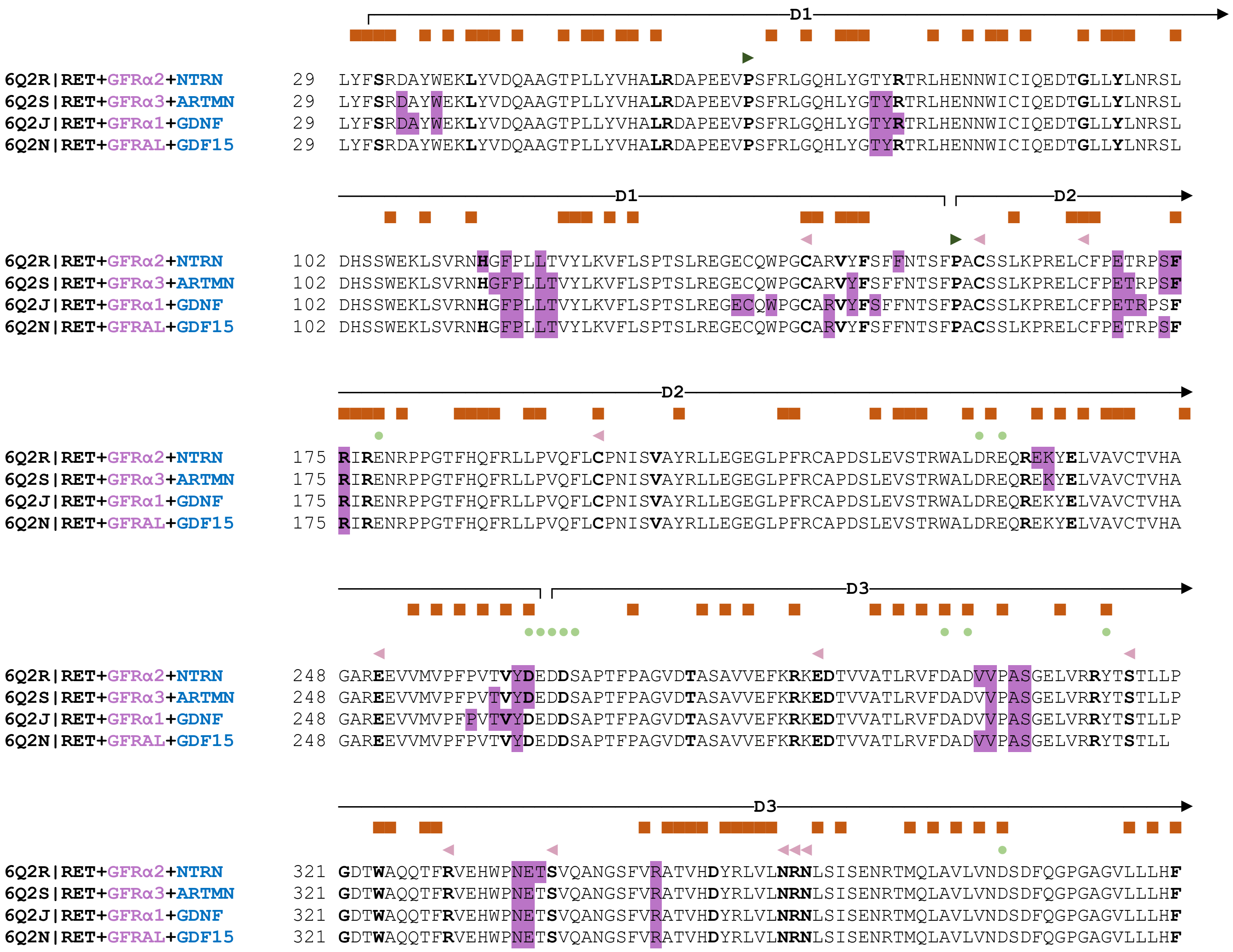
D1
6Q2R|RET+GFRα2+NTRN 29 LYFSRDAYWEKLYVDQAAGTPLLYVHALRDAPEEVPSFRLGQHLYGTYRTRLHENNWICIQEDTGLLYLNRSL
6Q2S|RET+GFRα3+ARTMN 29 LYFSRDAYWEKLYVDQAAGTPLLYVHALRDAPEEVPSFRLGQHLYGTYRTRLHENNWICIQEDTGLLYLNRSL
6Q2J|RET+GFRα1+GDNF 29 LYFSRDAYWEKLYVDQAAGTPLLYVHALRDAPEEVPSFRLGQHLYGTYRTRLHENNWICIQEDTGLLYLNRSL
6Q2N|RET+GFRAL+GDF15 29 LYFSRDAYWEKLYVDQAAGTPLLYVHALRDAPEEVPSFRLGQHLYGTYRTRLHENNWICIQEDTGLLYLNRSL
D1
D2
6Q2R|RET+GFRα2+NTRN 102 DHSSWEKLSVRNHGFPLLTVYLKVFLSPTSLREGECQWPGCARVYFSFFNTSFPACSSLKPRELCFPETRPSF
6Q2S|RET+GFRα3+ARTMN 102 DHSSWEKLSVRNHGFPLLTVYLKVFLSPTSLREGECQWPGCARVYFSFFNTSFPACSSLKPRELCFPETRPSF
6Q2J|RET+GFRα1+GDNF 102 DHSSWEKLSVRNHGFPLLTVYLKVFLSPTSLREGECQWPGCARVYFSFFNTSFPACSSLKPRELCFPETRPSF
6Q2N|RET+GFRAL+GDF15 102 DHSSWEKLSVRNHGFPLLTVYLKVFLSPTSLREGECQWPGCARVYFSFFNTSFPACSSLKPRELCFPETRPSF
D2
6Q2R|RET+GFRα2+NTRN 175 RIRENRPPGTFHQFRLLPVQFLCPNISVAYRLLEGEGLPFRCAPDSLEVSTRWALDREQREKYELVAVCTVHA
6Q2S|RET+GFRα3+ARTMN 175 RIRENRPPGTFHQFRLLPVQFLCPNISVAYRLLEGEGLPFRCAPDSLEVSTRWALDREQREKYELVAVCTVHA
6Q2J|RET+GFRα1+GDNF 175 RIRENRPPGTFHQFRLLPVQFLCPNISVAYRLLEGEGLPFRCAPDSLEVSTRWALDREQREKYELVAVCTVHA
6Q2N|RET+GFRAL+GDF15 175 RIRENRPPGTFHQFRLLPVQFLCPNISVAYRLLEGEGLPFRCAPDSLEVSTRWALDREQREKYELVAVCTVHA
D3
6Q2R|RET+GFRα2+NTRN 248 GAREEVVMVPFPVTVYDEDDSAPTFPAGVDTASAVVEFKRKEDTVVATLRVFDADVVPASGELVRRYTSTLLP
6Q2S|RET+GFRα3+ARTMN 248 GAREEVVMVPFPVTVYDEDDSAPTFPAGVDTASAVVEFKRKEDTVVATLRVFDADVVPASGELVRRYTSTLLP
6Q2J|RET+GFRα1+GDNF 248 GAREEVVMVPFPVTVYDEDDSAPTFPAGVDTASAVVEFKRKEDTVVATLRVFDADVVPASGELVRRYTSTLLP
6Q2N|RET+GFRAL+GDF15 248 GAREEVVMVPFPVTVYDEDDSAPTFPAGVDTASAVVEFKRKEDTVVATLRVFDADVVPASGELVRRYTSTLL
D3
6Q2R|RET+GFRα2+NTRN 321 GDTWAQQTFRVEHWPNETSVQANGSFVRATVHDYRLVLNRNLSISENRTMQLAVLVNDSDFQGPGAGVLLLHF
6Q2S|RET+GFRα3+ARTMN 321 GDTWAQQTFRVEHWPNETSVQANGSFVRATVHDYRLVLNRNLSISENRTMQLAVLVNDSDFQGPGAGVLLLHF
6Q2J|RET+GFRα1+GDNF 321 GDTWAQQTFRVEHWPNETSVQANGSFVRATVHDYRLVLNRNLSISENRTMQLAVLVNDSDFQGPGAGVLLLHF
6Q2N|RET+GFRAL+GDF15 321 GDTWAQQTFRVEHWPNETSVQANGSFVRATVHDYRLVLNRNLSISENRTMQLAVLVNDSDFQGPGAGVLLLHF

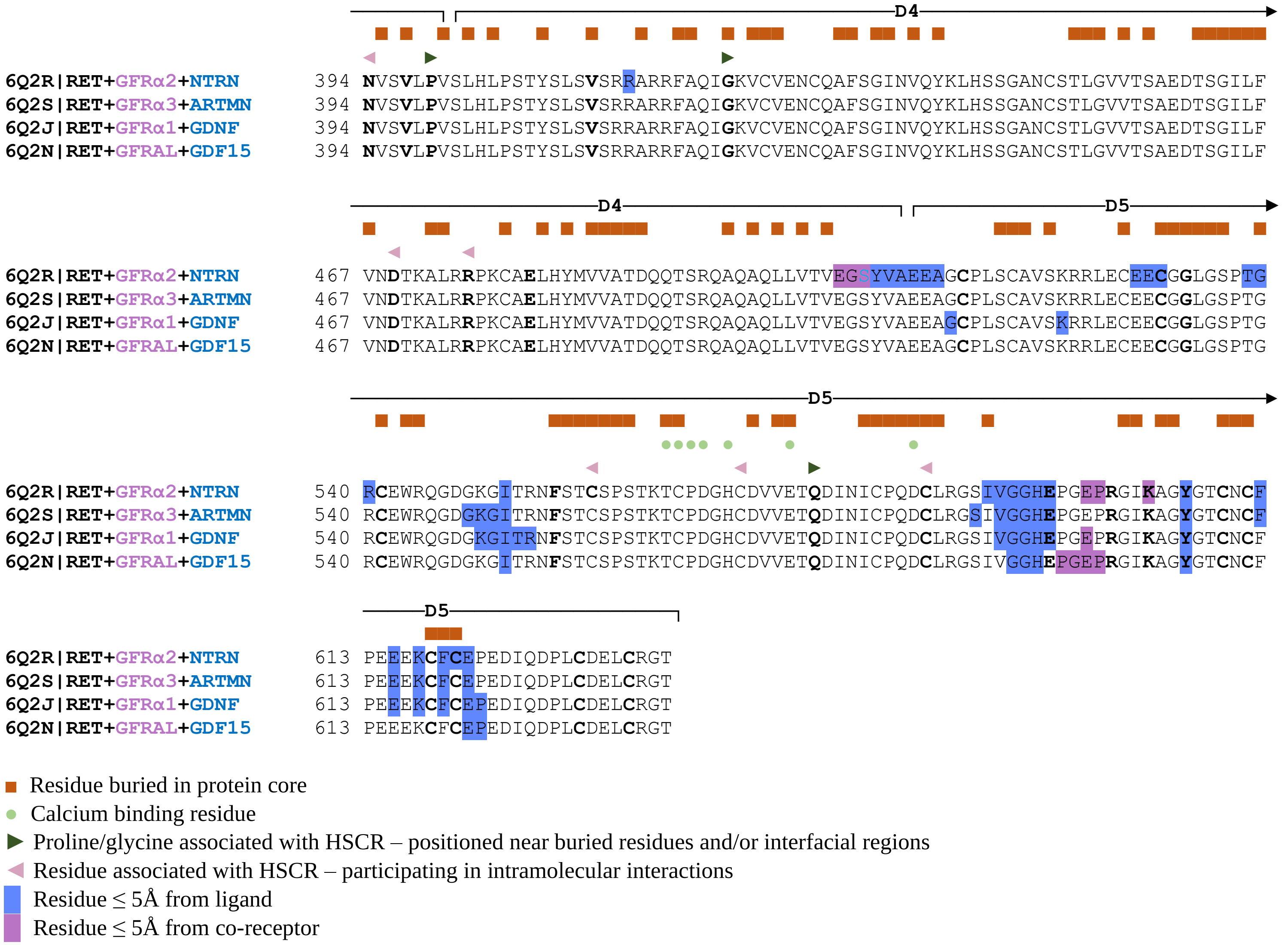


**Fig S1. Residues in RET structures that may affect protein tertiary structure or intermolecular interactions.** Residues that can affect tertiary structure, participate in intramolecular interactions, or in protein-protein interactions with ligands or co-receptors are annotated above the alignment according to the key. RET positions associated with MEN2A or HSCR diseases are marked in bold.

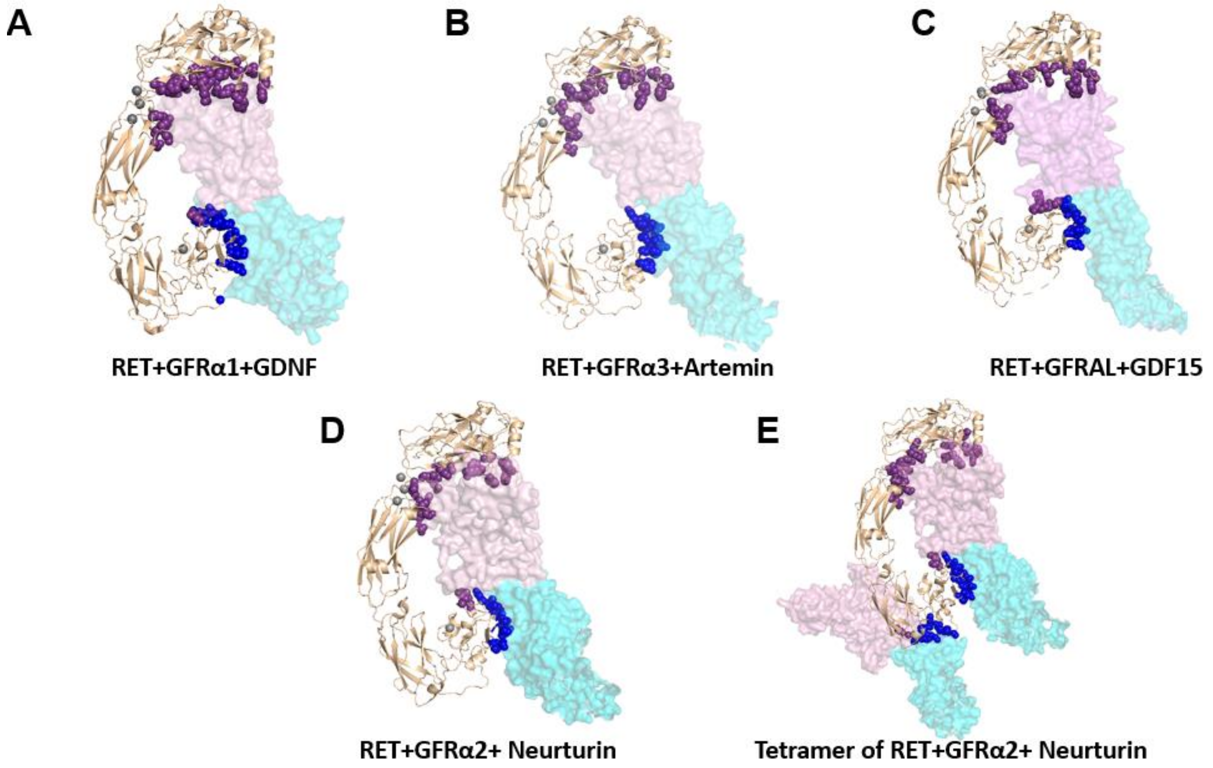


**Fig S2. Residues in the extracellular region of RET ≤ 5Å from ligands or co-receptors. A.** Residues in RET (wheat ribbon) ≤ 5Å of the GFRα1 co-receptor (pink molecular surface) or the GDNF ligand (cyan molecular surface) in PDB ID 6Q2J, shown as spheres and colored blue and purple, respectively. **B.** Residues in RET ≤ 5Å of the GFRα3 co-receptor (pink molecular surface) or the Artemin ligand (cyan molecular surface) in PDB ID 6Q2S, shown as in A. **C.** Residues in RET ≤ 5Å of the GFRAL co-receptor (pink molecular surface) or the GDF15 ligand (cyan molecular surface) in PDB ID 6Q2N, shown as in A. **D.** Residues in RET ≤ 5Å of the GFRα2 co-receptor (pink molecular surface) or the Neurturin ligand (cyan molecular surface) in PDB ID 6Q2O, shown as in A. **E.** Residues in RET ≤ 5Å of the two GFRα2 co-receptors (pink molecular surface) or the two Neurturin ligands (cyan molecular surface) in the RET tetramer complex in PDB ID 6Q2R, shown as in A.
